# Identification of Key Regulators in Pancreatic Ductal Adenocarcinoma using Network theoretical Approach

**DOI:** 10.1101/2024.03.12.584603

**Authors:** Kankana Bhattacharjee, Aryya Ghosh

## Abstract

Pancreatic Ductal Adenocarcinoma (PDAC) is a devastating disease with poor clinical outcomes, which is mainly because of delayed disease detection, resistance to chemotherapy, and lack of specific targeted therapies. The disease’s development involves complex interactions among immunological, genetic, and environmental factors, yet its molecular mechanism remains elusive. A major challenge in understanding PDAC etiology lies in unraveling the genetic profiling that governs the PDAC network. To address this, we examined the gene expression profile of PDAC and compared it with that of healthy controls, identifying differentially expressed genes (DEGs). These DEGs formed the basis for constructing the PDAC protein interaction network, and their network topological properties were calculated. It was found that the PDAC network self-organizes into a scale-free fractal state with weakly hierarchical organization. Newman and Girvan’s algorithm (leading eigenvector (LEV) method) of community detection enumerated four communities leading to at least one motif defined by G (3,3). Our analysis revealed 33 key regulators were predominantly enriched in neuroactive ligand-receptor interaction, Cell adhesion molecules, Leukocyte transendothelial migration pathways; positive regulation of cell proliferation, positive regulation of protein kinase B signaling biological functions; G-protein beta-subunit binding, receptor binding molecular functions etc. Transcription Factor and mi-RNA of the key regulators were obtained. Recognizing the therapeutic potential and biomarker significance of PDAC Key regulators, we also identified approved drugs for specific genes. However, it is imperative to subject Key regulators to experimental validation to establish their efficacy in the context of PDAC.

## Introduction

Pancreatic ductal adenocarcinoma (PDAC) is a common type of cancer originating from the pancreatic glands and is characterized by a rapidly progressive course and a dismal prognosis (1). Current research is actively exploring combination strategies that aim at various signaling pathways crucial for the growth and spread of tumors, particularly in pancreatic cancer. These endeavors hold the potential to significantly alter the approach to treating this disease (2). This is a highly detrimental disease with dismal clinical outcomes, primarily attributable to delayed disease detection, chemotherapy resistance, and absence of specific targeted therapies. The identification of novel therapeutic targets and/or early biomarkers for the disease has the potential to significantly improve the clinical management of PDAC and extend the lives of patients. To develop successful drug therapies, a deeper understanding of the molecular mechanisms underlying drug targeting is essential. A connectivity Map is a platform that provides information on the signaling pathways activated by a particular drug and can serve as a valuable resource for drug development. A promising strategy for treating PDAC involves exploiting aberrant metabolic processes in cancer cells, particularly PDAC cells. Cancer cells alter their metabolic pathways, a process regulated by complex and poorly defined interplay between intrinsic and extrinsic factors.

The incidence of PDAC is rising, and it is now one of the leading causes of cancer-related deaths in developed countries (3). Current treatment paradigms for PDAC, including targeted therapy, have shown limited success in improving survival outcomes (4). The molecular heterogeneity of PDAC and its complex tumor microenvironment contribute to its resistance to conventional treatments and immunotherapies (5) . Retinoids, such as retinoic acid (RA), have shown potential in maintaining normal pancreatic functions and have been explored as a therapeutic option for PDAC (6). The use of predictive molecular markers and cancer gene panel testing may help in selecting personalized therapies for PDAC patients (7). Accurate diagnosis of PDAC is crucial, as it can be easily misdiagnosed as other pancreatic neoplasms, such as acinar cell carcinoma (ACC) or neuroendocrine tumor (PNET). Further research is needed to overcome the challenges in PDAC management and improve patient outcomes. Recent research is exploring a wide range of novel therapeutic targets for PDAC, including genomic alterations, tumor microenvironment, and tumor metabolism (8). Advancements in tumor genome sequencing technologies are rapidly occurring, opening avenues for tailored and targeted therapies for PDAC (9). Immunotherapy with immune checkpoint inhibitors has shown promise in PDAC, particularly in tumors harboring mismatch repair deficiency (dMMR) and high microsatellite instability (MSI-H) (10). The presence of a dense fibrotic stroma in PDAC creates a physical barrier around the cancer cells, hindering drug delivery and promoting tumor growth and treatment resistance (11). Advances in surgical approaches, immunotherapeutic approaches, and targeted therapies are being explored to overcome the treatment refractory nature of PDAC and improve patient outcomes (12). The tumor microenvironment (TME) plays a crucial role in PDAC development, progression, and treatment resistance (13). Immunotherapy, including checkpoint inhibition and immune-based therapies, has shown promise in PDAC treatment (14). Additionally, targeted therapies that focus on specific signaling pathways and components of the TME, such as fibro-blasts and cancer-associated fibroblasts, are being explored .

Pancreatic ductal adenocarcinoma (PDAC) is an aggressive and deadly cancer with limited treatment options. Current standard of care treatments for advanced PDAC include systemic chemotherapy regimens such as gemcitabine/nab-paclitaxel and FOLFIRINOX, which have improved clinical outcomes (15). However, the 5-year survival rate for PDAC remains low, highlighting the need for new therapies (16). The tumor microenvironment (TME) of PDAC plays a significant role in tumorigenesis and may contain promising novel targets for therapy (17). Adjuvant chemotherapy has been shown to offer a survival advantage over surgery alone in resected PDAC (18). However, the addition of chemoradiation therapy (CRT) to chemotherapy does not provide a survival advantage and is not recommended (19). Surgical resection is the first choice for treatment of pancreatic acinar cell carcinoma (PACC), but there is no standard treatment option for inoperable disease . Understanding the risk factors for PDAC can help in screening and counseling patients for lifestyle modifications .

A disorder typically arises from disruptions within the intricate internal web of connections (20) among genes that serve related functions, rather than from the abnormality of a single gene. This resulted in the adoption of a systemic approach to biological issues, founded on the principle that comprehending the involvement of different genes/proteins is essential (21) in disease initiation and progression . It is necessary to consider the entire network of interactions within a living system (22). In this context, it is important to employ Network Medicine (23), as this method aids in the investigation of disease pathways and modules linked to complex illnesses.

In this study, the protein interaction maps are analyzed through graph/network theory to get insights about the theoretical aspect of complex networks (24). According to the graph theory, analysis of the topological structure of a network (PPI network in our study) provides important information of the network (25) through which novel disease genes and pathways, biomarkers and drug targets for complex diseases can be identified (26).

Thus, the focus of our study is on the protein-protein interactions network/graph of PDAC, constructed from differentially expressed genes with an aim to understand the architectural principle of the network/graph (random, small world, scale-free or hierarchical). We further extended our study to the prediction of important key regulators of the network which have fundamental importance due to their activities and regulating mechanisms in the network (27). It is expected that the findings of this study will advance our understanding of the initiation and progression of PDAC, thereby, strengthening different therapeutic approaches for PDAC. This work was to establish an unbiased catalogue of change in (up and down regulation) gene expression for the PDAC samples. we bestowed 12 samples of PDAC analysis of gene-expression from normal and cancer patients. Finally, the comparative study of these conditions (PDAC Cancer and Normal) has affirmed the thirty-three (33) prominent genes. These are intricated in basic functioning of the cell like as in the rearrangement of the cytoskeleton, tissues development and activation of immune system. This work will be useful for the researchers to collect evidences against PDAC working in in-vitro.

## Materials and Methods

### Preprocessing and acquisition of dataset

The RNA-seq dataset GSE171485 (28) was downloaded from the NCBI GEO database [https://www.ncbi.nlm.nih.gov/].

### Data Analysis and visualization of Differentially Expressed Genes

The data analyses were conducted using R Studio (4.2.2) and R .The fold change method was employed to identify differentially expressed genes. The fold change for a gene is calculated by subtracting the intensities measured in two samples (control vs clinical groups). This value is referred to as the fold change. These raw values are typically log transformed (usually log2) (29). Another method to calculate the fold change ratio involves dividing the two measured intensities for a given gene in two samples. A change of at least two-fold (up or down regulated) considered to be significant.

### Screening of DEGs

We utilized the LIMMA package (30) to analyze the data and identify Differentially expressed genes (DEGs) by assessing gene expression values. This approach employs Linear Modeling (LM) (31) and Empirical Bayes (EB) (32) techniques to perform f-tests (33) and t-tests (34), while simultaneously reducing standard errors. The objective of this method is to produce results that are dependable and reproducible, thereby enhancing the precision and reliability of the statistical analyses. DEG analysis was conducted to compare gene expression levels between control and clinical groups. The R package “ggplot2” (35) was employed for data visualization purposes. The fold change criteria for each gene were established based on a Benjamini-Hochberg adjusted p-value threshold of 0.05 and a significance level of p ≤ 0.05.

### Construction of Protein-Protein Interaction Network of PDAC

The construction of the PDAC PPI (Protein-Protein Interaction) network was undertaken using the STRING database (The Search Tool for the Retrieval of Interacting Genes, (<http://string-db.org/>) with an interaction score threshold of > 0.4. This approach enables the exploration and analysis of protein-protein interactions, which can be either physical or functional associations. These associations are derived from text-mining of literature, co-expression analysis, genomics context-based predictions, computational predictions, and high-throughput experimental data, as well as the aggregation of previous knowledge from other databases. The network was subsequently visualized and analyzed using the Cytoscape software (version 3.6.1) (36).

### Gene ontology (GO) enrichment analysis

Gene ontology terms provide a controlled vocabulary that is divided into three categories: Molecular Function, Biological Process, and Cellular Location. To conduct a preliminary investigation into the functional differences of DEGs, they were submitted to DAVID (Database for Annotation, Visualization and Integrated Discovery), an online software (<http://david.abcc.ncifcrf.gov/home.jsp>), to enrich the set of DEGs with possible GO terms (37).

### Delineation of global Network topological properties

The structure of the network can be examined through its topological properties, which establish the connections between nodes and illustrate their interactions. The topology of the network is defined by the probability of degree distributions (*P*(*k*)), clustering coefficient (P(k)), and neighborhood connectivity (C_N_(k)), which exhibit a power law distribution. This adherence to the power law distribution indicates the presence or absence of scale-free properties in the network. The network constructed for PDAC using differentially expressed genes extracted from the RNA-seq dataset GSE171485 follows the power law distribution in its probability of degree distributions (P(k)), clustering coefficient (*C*(*k*)), and neighborhood connectivity (C_N_(k)). This is depicted in **Figure 4** and confirmed by the matrix representation of all properties (38).

The network’s behavior is characterized by equations (see eq 1, 2, 3) that reveal a hierarchical structure in synergy with the scale-free nature of the network. The power law distribution was fitted to the topological properties of the network using a standard statistical fitting procedure.

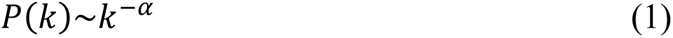

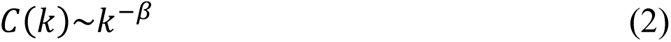

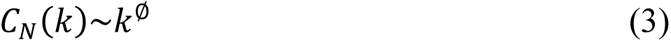

The positive value in phi of connectivity parameter shows assortative nature of the network. While, the negative value in alpha (α) of degree distribution shows availability of each node in the network. The negative value in beta of clustering parameter shows disassortative in the communication between the nodes in network (39).

### Communities finding: LEV method

To investigate the nature and topological properties of hierarchical networks at various levels of organization is beneficial for understanding network behavior and uncovering the organizing principles that govern them. There are several methods for detecting communities within these networks, one of which is the leading eigenvector (LEV) method. This method calculates the eigenvalue for each edge, giving greater importance to links rather than nodes. For this study, we applied the LEV detection method using the ’igraph’ (40) package in R. We used this code to detect modules in the main network, sub-modules within each module at different levels of organization, and so on, ultimately identifying motifs (i.e. 3 nodes and 3 edges) (41). Throughout this process, we adhered to the criterion of identifying any sub-module as a community if it contained at least one motif (defined by G (3,3)).

### TF/miRNAs screening

To identify miRNAs targeting key signature genes, we employed MIENTURNET (MicroRNA ENrichment TURned NETwork). MIENTURNET (42) is an interactive web application designed for micro-RNA target enrichment analysis, primarily relying on the TargetScan program for sequence-based miRNAs target predictions (43). By utilizing the MIENTURNET software, we successfully achieved significant functional enrichment of the predicted miRNAs.

### Drug Gene Interaction Analysis

To identify potential drugs for treating PDAC, we utilized the DGIdb web tool (44) a database of drug-gene interactions and druggable genes.

## Results

This work provides information in RNA-seq dataset GSE171485 on the structure and function of interacting genes. The results of the differential expression analysis, as shown in volcano plot (**Figure 1**), indicate that 772 DEGs were identified. Of these, 341 genes were found to be down-regulated and 431 were up-regulated, with the threshold cut-offs set at log2 |FC| >= 1, P − value < 0.05, and Padj ≤ 0.05.

**Figure 1.**
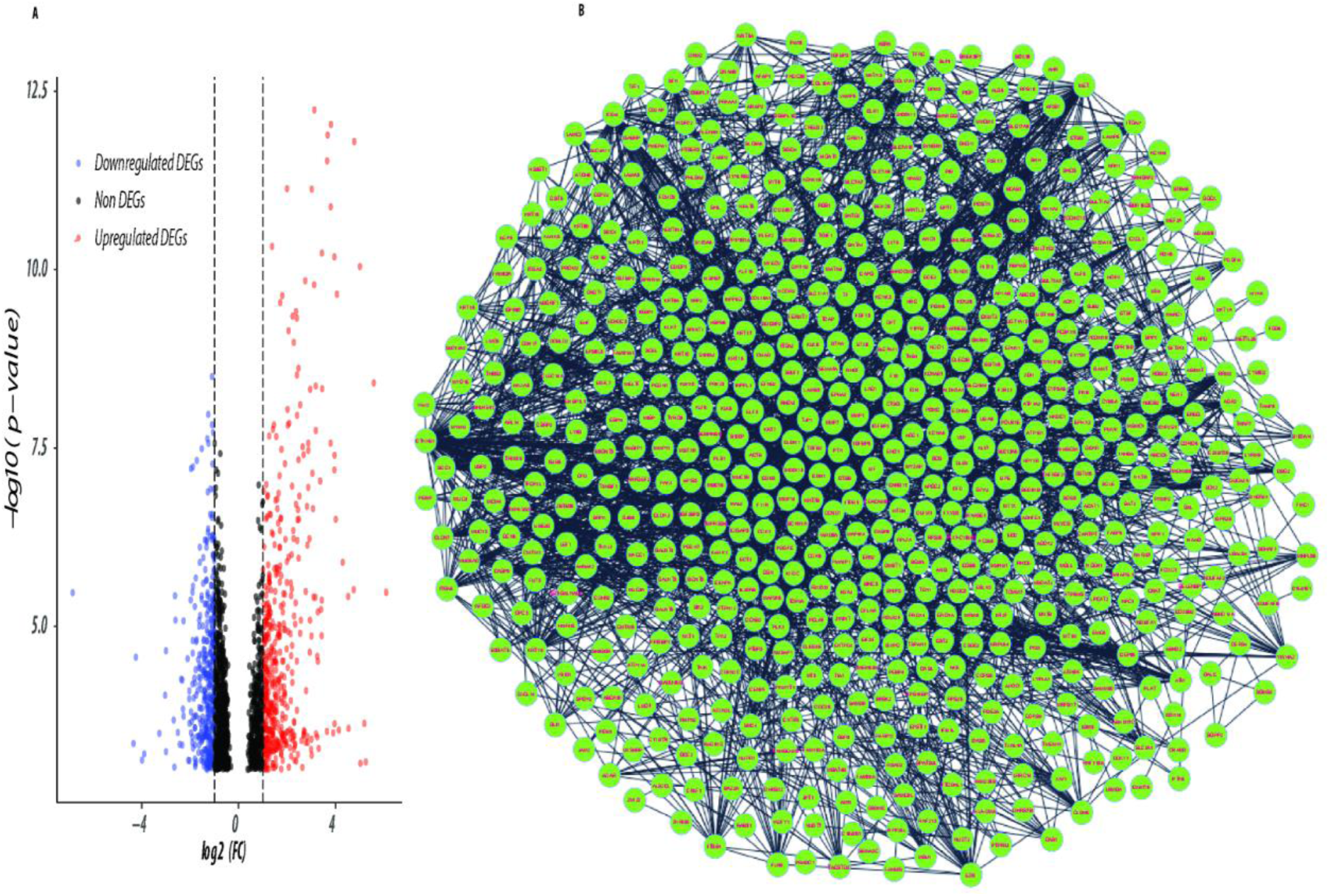
Figure showing (A) volcano plot showing up & down regulated expressed genes (B) Network of differentially expressed genes.

To get the idea of what could be the effect of DEGs can only be possible when we have some preliminary insight in to their individual function. As the term gene ontology (GO) enrichment gives us the opportunity to get the basic idea about any gene. GO enrichment analysis of significantly enriched DEGs between disease and control was categorized into biological process, cellular components and molecular function. Among these down-regulated DEGs, potassium ion transport, cell-cell signaling, neuropeptide signaling pathway were most significantly down-regulated biological process in disease (Figure 2A). While other important biological process associated with these down-regulated DEGs were muscle contraction, cellular response to zinc ion, response to hypoxia, negative regulation of cytosolic calcium ion concentration and positive regulation of apoptotic process. While most significantly down-regulated cellular components and molecular function associated with the DEGs were extracellular region, mitochondrion, potassium channel complex, endoplasmic reticulum, mitochondrion inner membrane and protein binding, methyltransferase activity, structural constituent of ribosome respectively (Figure 2B & 2C). KEGG pathways enrichment of down-regulated genes were enriched in metabolic pathway, peroxisome cAMP signaling pathway and chemical carcinogenesis-reactive oxygen species (Figure 2D).

**Figure 2.**
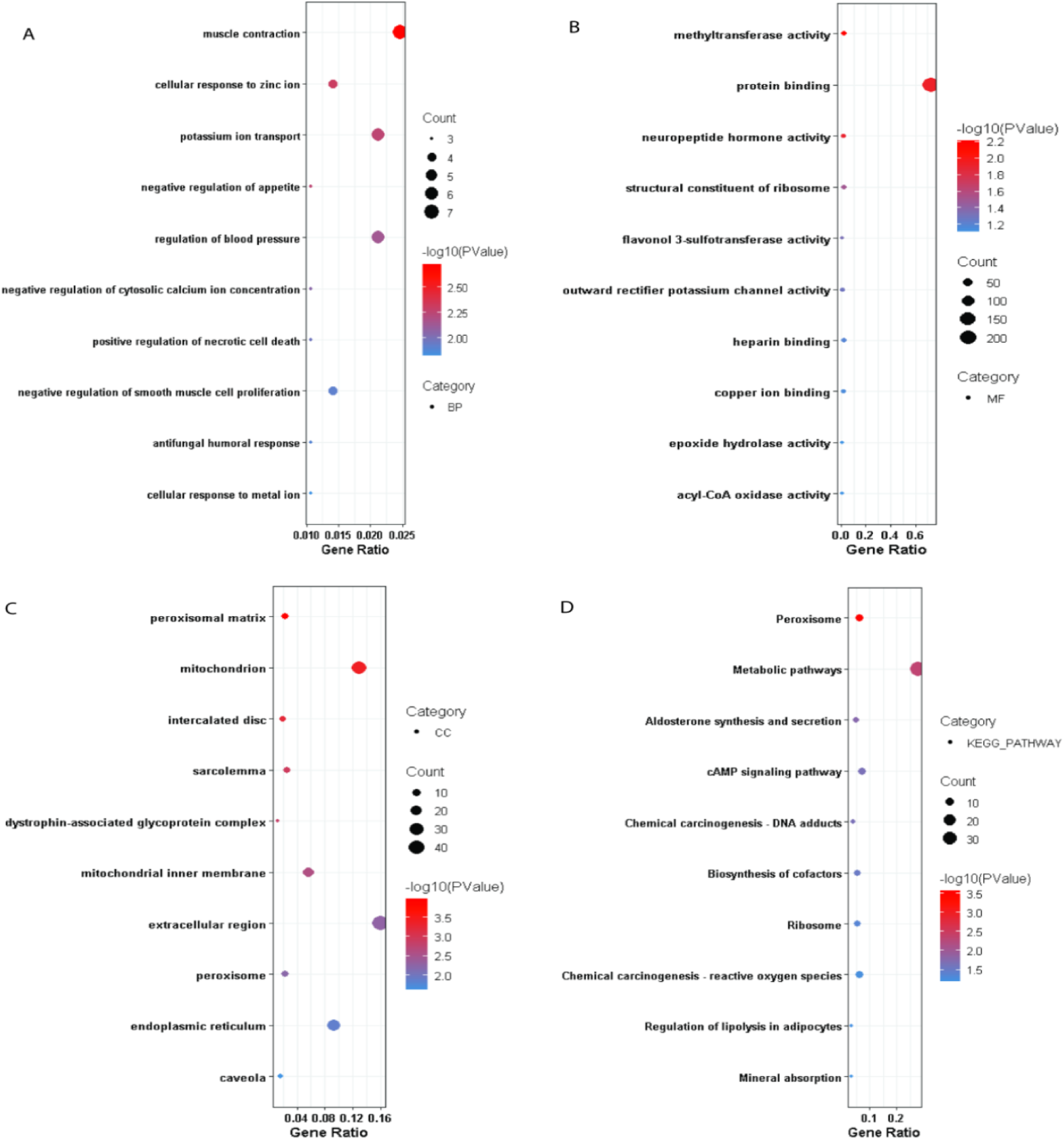
Gene Ontology analysis for down-regulated genes of pancreatic ductal adenocarcinoma.

Among the top 15 up-regulated DEGs, cell-cell adhesion, positive regulation of cell migration, extracellular matrix organization, tissue development, epithelial cell differentiation and integrin-mediated signaling pathway were most significantly up-regulated biological process in PDAC (Figure 3A). GO classification indicates the up-regulation of calcium ion binding, calcium-dependent protein binding, cadherin binding, virus receptor activity and insulin-like growth factor I binding related molecular function in PDAC (Figure 3B).

**Figure 3.**
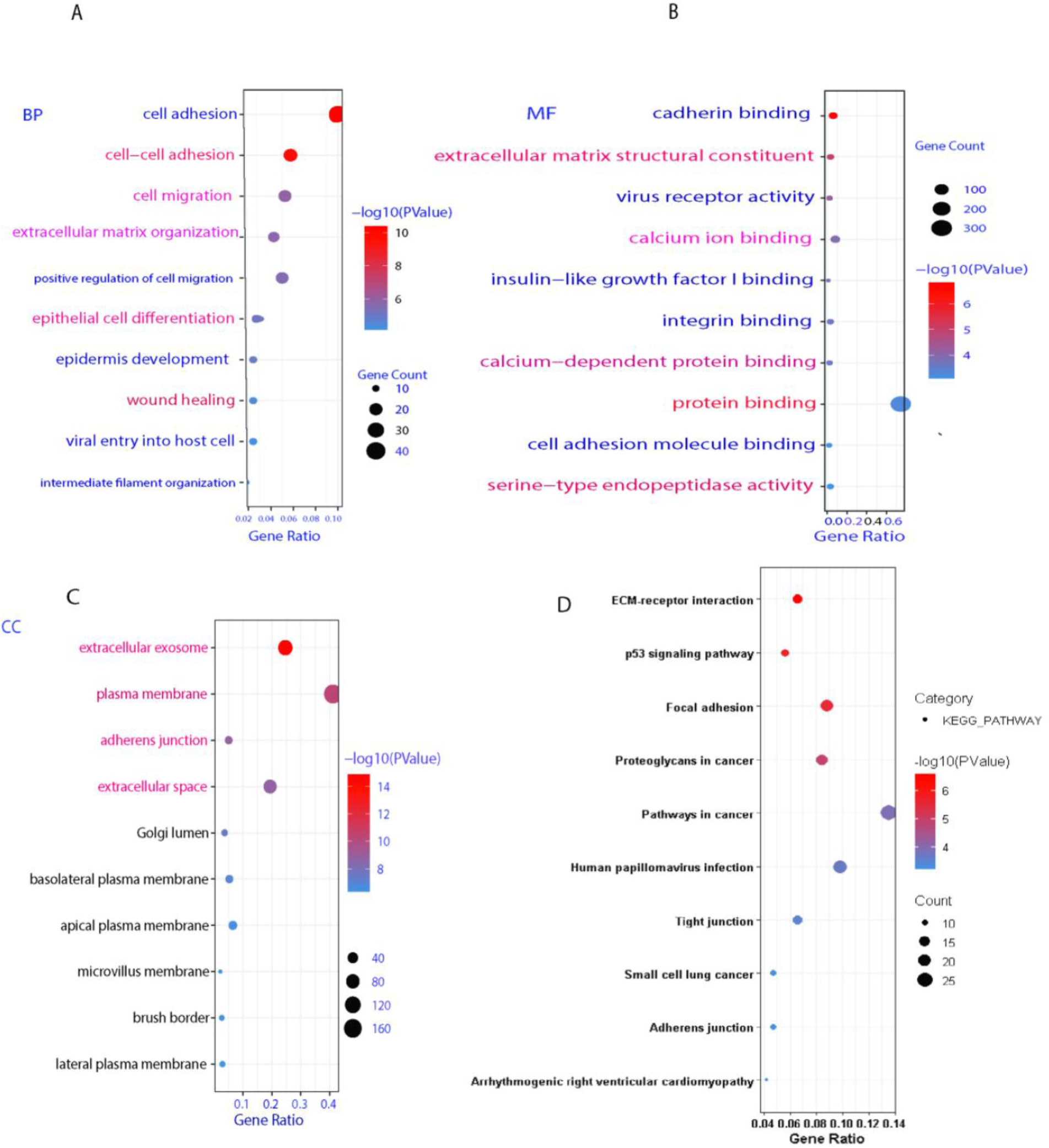
Gene Ontology analysis for up-regulated genes of pancreatic ductal adenocarcinoma.

While most significantly up-regulated cellular components were plasma membrane, extracellular exosome, and perinuclear region of cytoplasm Figure as shown in (Figure 3C). KEGG pathways were enriched in ECM-receptor interaction, p53 signaling pathway, pathways in cancer and proteoglycan in cancer (Figure 3D).

### PDAC Network Architecture Reveals Hierarchical Scale-free Features

To gain insights into the PPI network of PDAC’s structural features, we analyzed its topology, specifically the probability of degree distribution *P*(*k*), clustering coefficient *C*(*k*) neighborhood connectivity C_N_(k), and centrality measurements. We employed the statistical fitting technique proposed by Clauset et al. (45) to verify that the graph’s architecture follows power-law behavior. Our results indicate that all statistical p-values, calculated against 2500 random samplings, are greater than the critical value of 0.1, and the goodness of fits are less than or equal to 0.35. The data points of all the topological parameters fit power law when plotted against the degree k of the PDAC network.

The values of the power-law exponents for each of the topological properties of the complete network were calculated:

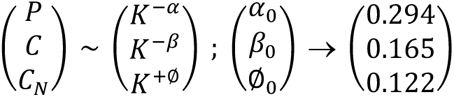

The negative values of α = 0.294; (α < 2) and β = 0.165; (β < 1) suggest the hierarchical nature of the PD network, indicating the existence of well-defined successive interconnected communities with sparsely distributed hubs in the network. The values of the exponents of *P*(*k*), *C*(*k*), and *C_N_*(*k*)(∅Ø = 0.122; (+∅Ø ≤ 0.5)) suggest that the network, though not strongly hierarchical, falls into the category of a weak hierarchical scale-free network. The negative value of β indicates that as k increases, C(*k*) decreases, suggesting that nodes with a high degree have a low tendency to cluster, further indicating a hierarchy of hubs, in which the most densely connected hub is linked to a small fraction of all other nodes. The power-law distribution observed in *P*(*k*) is indicative of the scale-free nature of the small-world network, with the negative value of α indicating that a small number of nodes possess a high degree while the majority of nodes have a low degree, which is consistent with the network’s scale-free behavior. The positive value of ∅Ø suggests that the network exhibits assortative mixing, where edges predominantly connect heavily connected nodes, regulating the system’s behavior. Further,

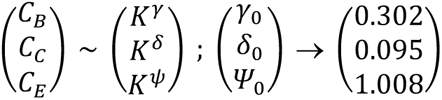

The positive values of the exponents γ, δ, and ψ of the three distributions CB(k), Cc(k) and CE(k) respectively, indicate that the network exhibits hierarchical scale-free or fractal features. These positive values signify that CB(k), Cc(k) and CE(k) increase as the degree k increases when plotted against it (Figure 4 C, D &F). The increasing value of CB(k) as k increases suggests that nodes with a high degree have high CB(k), indicating that these larger hubs have a major influence on the information transmission in the network compared to nodes with a low degree. Similarly, the direct proportionality between Cc(k) and k suggests that high-degree nodes are quick spreaders of information in the network, indicating their high Cc(k). The positive value of ψ indicates that nodes with a high degree have high CE(k) as well, suggesting their influence in the network due to their ability to spread information. The positive value of ψ also signifies the connectedness between high-degree nodes, which is in agreement with the assortative mixing in the network. Through a meticulous study of these topological properties, it was found that the PDAC network self-organizes into a scale-free fractal state with weakly hierarchical organization.

**Figure 4:**
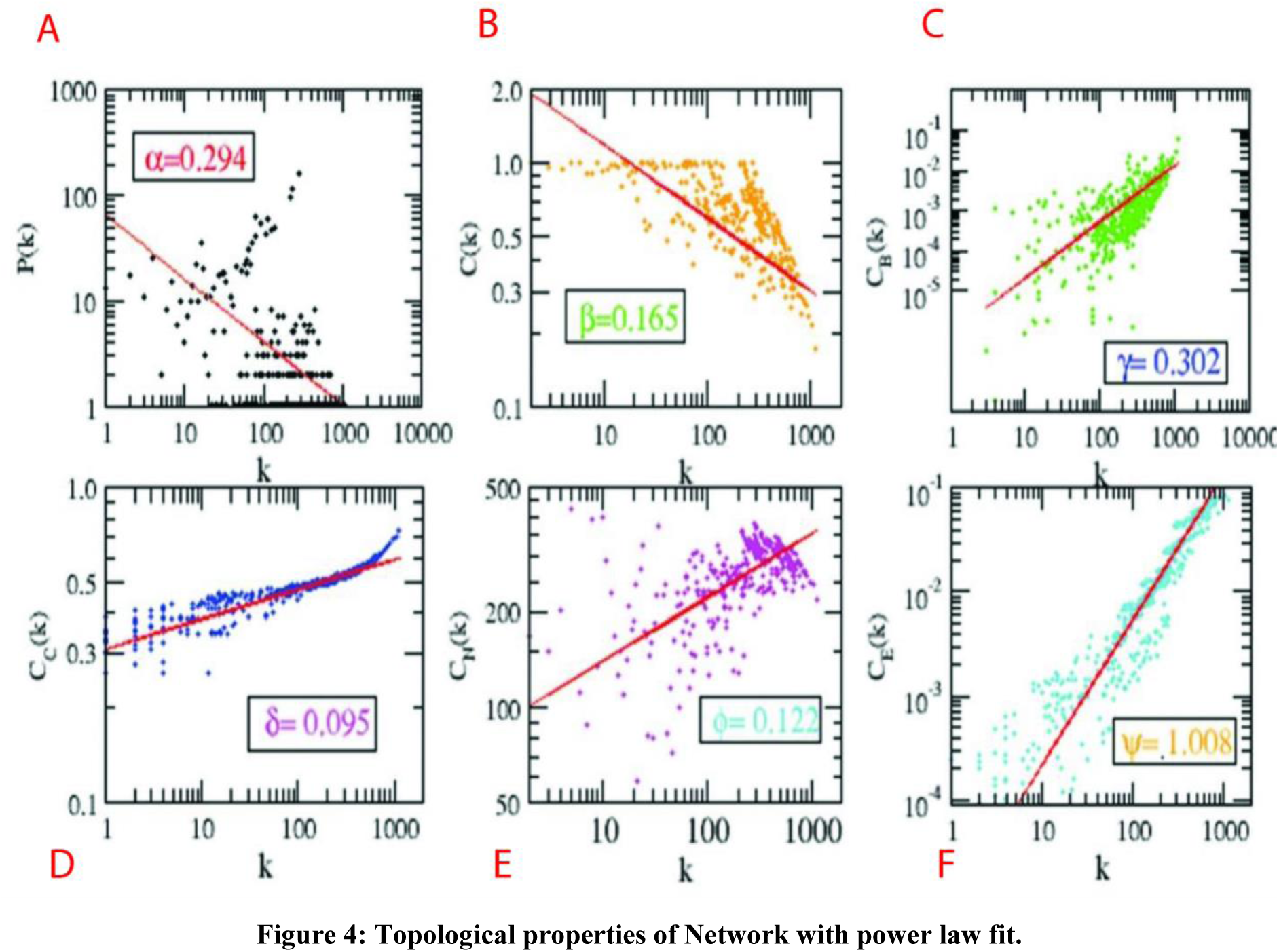
Topological properties of Network with power law fit.

### Key Regulators Uncovered through Clustering and Tracing

Owing to the significance of regulator genes(RGs) as functional bottlenecks in the initiation and progression of a disease by regulating the expression of a plethora of downstream effector genes (46), we identified the most potent RGs of the PDAC network. Newman and Girvan’s algorithm helped us untangle the PDAC network and the network was observed to be organized in five hierarchical levels using this algorithm (Figure 5). After tracing of the G (3,3) triangular structure genes from top to bottom organization through these levels of hierarchy, 33 genes were revealed to be the RGs of the PDAC network, the criterion being their presence at every topological level (Figure 5). This agrees with the definition of RGs, according to which, RGs are the genes/proteins which are deeply rooted from top to bottom organization of the network. These RGs are the backbone in maintaining a network’s stability as they capacitate the network to combat any unacceptable alterations in it.

**Figure 5.**
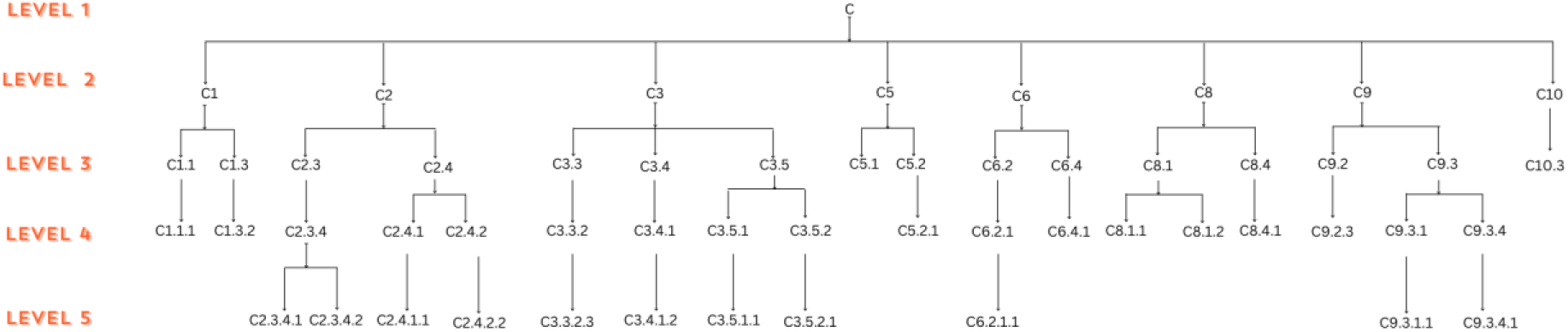
Tracking down the presence of the fundamental genes within different Modules at different level of the network.

*RNF213, EPSTI1* and XAF1 separated their way from the rest of the RGs from the first level itself and then took the path hand in hand till the motif level. Whereas, *GAL, VIP*, *GNRH1,NMU, VIPR2*, and GNG11 RGs all moved into a different sub-module of the first level and took path at the motif level. Genes *MMP1, CTSG, F2RL1, F10, PLAT, F2R, PDGFA, FGF13, PTN, APOC2, SDC1,* and *SDC4* moved into same module at level 2. Further, genes *MMP1, CTSG, F2RL1, F10, PLAT,* and *F2R* makes sub-module and belongs to same sub-module till level 4 and separated at level 5 as triangular motif structure (**Table 1**)

**Table 1:** Network breaking mechanism to understand the gene names and their sub-modules.

| Network decomposition outline | Gene names in motifs |
| --- | --- |
| C→C2→C2.3→C2.3.4→C2.3.4.1 | *F10*, *PLAT*, *F2R* |
| C→C2→C2.3→C2.3.4→C2.3.4.2 | *MMP1*, *CTSG*, *F2RL1* |
| C→C2→C2.4→C2.4.1→C2.4.1.1 | *PDGFA*, *FGF13*, *PTN* |
| C→C2→C2.4→C2.4.2→C2.4.2.2 | *APOC2*, *SDC1*, *SDC4* |
| C→C3→C3.3→C3.3.2→C3.3.2.3 | *AP2B1*, *AP1S3*, *DNM2* |
| C→C3→C3.4→C3.4.1→C3.4.1.2 | *EGFL7*, *RASIP1*, *SOX18* |
| C→C3→C3.5→C3.5.1→C3.5.1.1 | *CLDN2*, *CLDN7*, *CDH3* |
| C→C3→C3.5→C3.5.2→C3.5.2.1 | *CLDN5*, *TJP1*, *PCDH1* |
| C→C6→C6.2→C6.2.1→C6.2.1.1 | *RNF213*, *EPSTI1*, *XAF1* |
| C→C9→C9.3→C9.3.1→C9.3.1.1 | *VIP*, *GAL*, *GNRH1* |
| C→C9→C9.3→C9.3.4→C9.3.4.1 | *NMU*, *VIPR2*, *GNG11* |

In the third level, *PDGFA, FGF13, PTN, APOC2, SDC1,* and *SDC4* moved into the same sub-module and got separated at level four into two sub-modules. Afterward, these RGs moved separately till they reached the motif level i.e., the 5th level. Genes *AP2B1*, *AP1S3 DNM2*, *EGFL7*, *RASIP1, SOX18, CLDN2*, *CDH3, TJP1, CLDN5* and *PCDH1* clustered in same sub-module at level 2 and got separated at levels 3, 4 and reached motif levels (Figure 6).

**Figure 6.**
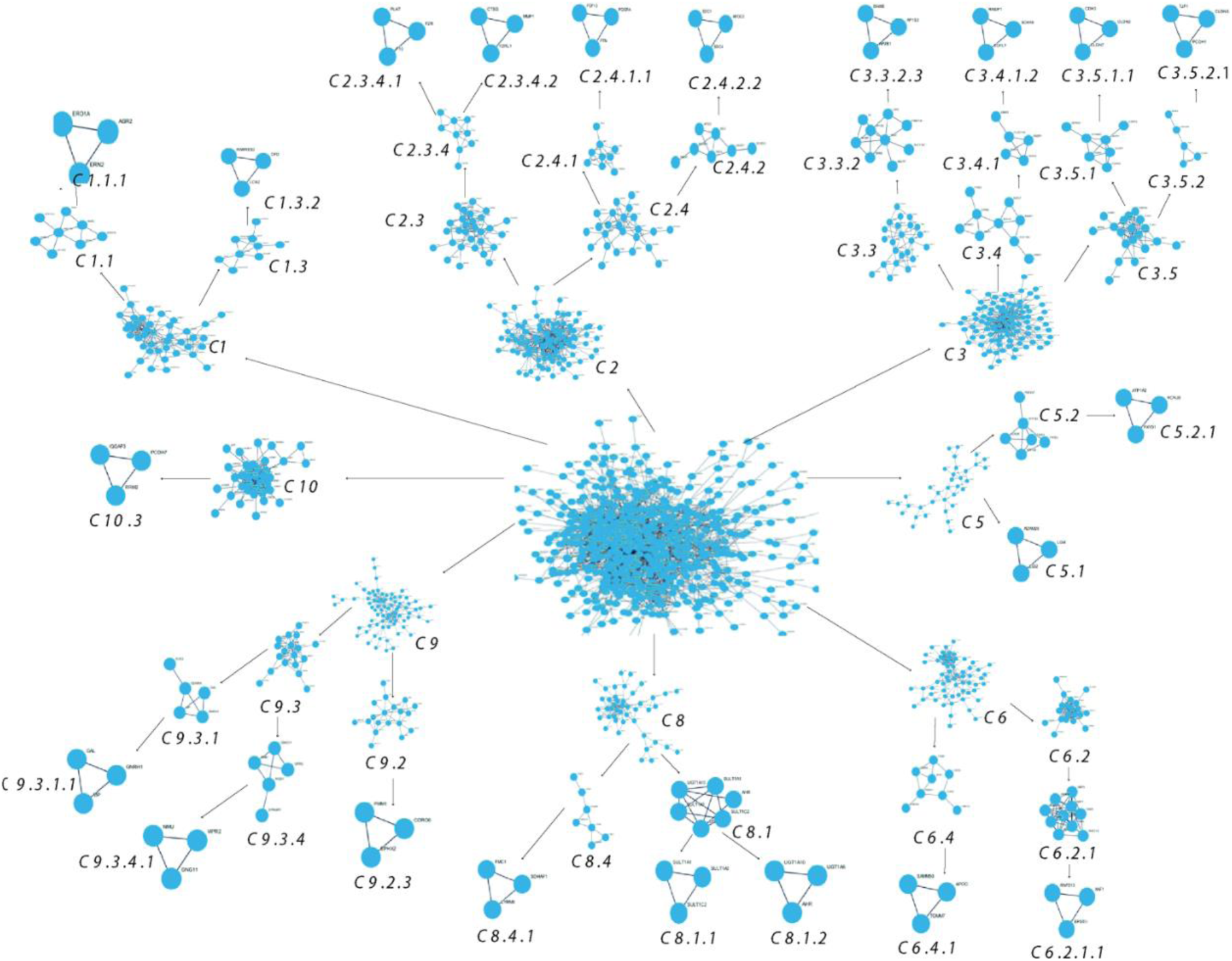
Detecting communities within PDAC networks using leading eigenvector (LEV) method.

### Emergence of Low Degree Node Accompanied by High Degree Node as Key Regulators

In a network, when a node’s degree is low, the node gains what strength it has from its neighbors and thus the influence it has over the network is a function of its neighbouring degree. Whereas for high degree nodes, the strength of the nodes comes from their large number of connections rather than their neighbouring degree (47). In addition, a low degree bridge node, connecting two high degree nodes, is very important in a network despite its lower degree (48). Thus, a node’s degree is not the sole determinant of its essentiality, rather, it depends on the topological position of that node. This is reflected in our results, where, *F2RL1, F10, APOC2, CLDN2, PCDH1* which has quite a low degree (4,4,8,11,3) respectively in the primary network, found out to be RGs based on its ability to make it to the last level of the organization. *F2RL1* formed a motif in the last level of the organization with another KR, *MMP1*, which has a fairly high degree (36) in the primary network. Gene *F10* formed a motif with *PLAT* having degree *17, APOC2* formed motif with *SDC1* having degree *40* and *PCDH1* formed motif with *TJP1* which fairly high degree is (44) in primary network. This shows that a low degree node i.e., *F2RL1, F10, APOC2, CLDN2,* and *PCDH1* are also an important information propagator in the network, contributing to serving as the backbone of the network and functioning (**Table 2**).

**Table 2:** Topological statistical properties of low degree node aaccompanied by high degree node as key regulators.

| S.N. | Gene name | Degree | Betweenness Centrality | Closeness Centrality | Clustering Coefficient |
| --- | --- | --- | --- | --- | --- |
| 1 | *F2RL1* | 4 | 0.0007 | 0.2791 | 0.6667 |
| 2 | *F10* | 4 | 0.0008 | 0.3041 | 0.1666 |
| 3 | *APOC2* | 8 | 0.0066 | 0.3030 | 0.2142 |
| 4 | *CLDN2* | 11 | 0.0009 | 0.3351 | 0.8727 |
| 5 | *PCDH1* | 3 | 0.0001 | 0.2835 | 0.3334 |
| 6 | *MMP1* | 36 | 0.0241 | 0.3742 | 0.2730 |
| 7 | *PLAT* | 17 | 0.0091 | 0.3517 | 0.2332 |
| 8 | *SDC1* | 40 | 0.0218 | 0.3735 | 0.2846 |
| 9 | *TJP1* | 44 | 0.0275 | 0.3796 | 0.1987 |

Furthermore, gene-drug interaction from DGIdb were retrieved and listed in Table 3.

**Table 3:** Gene-drug interaction of the key regulators:

| S.N. | Gene | Drug |
| --- | --- | --- |
| 1 | *F10* | EDOXABAN, IDRAPARINUX SODIUM, APIXABAN, RIVAROXABAN, HEPARIN, EDOXABAN TOSYLATE, EMICIZUMAB, BETRIXABAN, LETAXABAN, MENADIONE, OTAMIXABAN, IDRAPARINUX, TANOGITRAN, MELAGATRAN, TINZAPARIN, DANAPAROID, CHEMBL1271162, SEMULOPARIN, MANGIFERIN HEPTASULFATE, THROMBIN, TRIMETHOPRIM/SULFADOXINE |
| 2 | *PLAT* | AMINOCAPROIC ACID, ATORVASTATIN, MELPHALAN, EPOETIN BETA, RALOXIFENE, NAPROXEN, UROKINASE, BORTEZOMIB |
| 3 | *F2R* | RIGOSERTIB SODIUM, ATOPAXAR, VORAPAXAR SULFATE, VORAPAXAR, ATROPINE, WORTMANNIN, ARGATROBAN, |
|  |  | BLEOMYCIN, LEPIRUDIN, THALIDOMIDE, MORPHINE, ALCOHOL, ASPIRIN, THROMBIN, DALTEPARIN, RUSALATIDE |
| 4 | *MMP1* | DOXYCYCLINE CALCIUM, APRATASTAT, DOXYCYCLINE, DOXYCYCLINE HYCLATE, CIPEMASTAT, MARIMASTAT, PRINOMASTAT, LEUPROLIDE ACETATE, SIROLIMUS, COLLAGENASE CLOSTRIDIUM HISTOLYTICUM, MEDROXYPROGESTERONE ACETATE, LAMIVUDINE, LEFLUNOMIDE, HYDROCORTISONE, PENTOSAN POLYSULFATE SODIUM, TRIAMCINOLONE, RIBAVIRIN |
| 5 | *CTSG* | MANNITOL, CHEMBL374027 |
| 6 | *F2RL1* | CHEMBL493076, CHEMBL494502, ROXITHROMYCIN, MINOCYCLINE, TETRACYCLINE, ERYTHROMYCIN, CLARITHROMYCIN, DOXYCYCLINE |
| 7 | *PDGFA* | SUNITINIB, SQUALAMINE |
| 8 | *SDC1* | HEPARIN, INDATUXIMAB RAVTANSINE |
| 9 | *EGFL7* | PARSATUZUMAB |
| 10 | *CDH3* | PF-06671008 |
| 11 | *TJP1* | RISPERIDONE, GENISTEIN, ALCOHOL, DEXAMETHASONE |
| 12 | *VIP* | DIGOXIN, LISINOPRIL, OMEPRAZOLE, RIBAVIRIN, ANDROSTANOLONE, AZASERINE, FLUTAMIDE |
| 13 | *GAL* | LIOTHYRONINE SODIUM, HYDROCORTISONE, PRASTERONE, DIACETYLMORPHINE, MIRTAZAPINE |
| 14 | *GNRH1* | RESERPINE, RALOXIFENE, GOSERELIN, AMINOGLUTETHIMIDE, ZALCITABINE, DAPSONE, CAPTOPRIL, CHEMBL208519, LITHIUM, LEUPROLIDE, DITIOCARB |

Next, transcription factors were identified with TargetScan and listed in Table 4.

**Table 4:** Transcription factor of Key regulator genes.

| **S. N.** | **Key TF** | **Description** | **P-value** | **Key regulator Genes** |
| --- | --- | --- | --- | --- |
| 1 | NF1 | neurofibromin 1 | 0.000266 | *GNRH1, PLAT* |
| 2 | SP3 | Sp3 transcription factor | 0.001 | *F2R, PLAT, SOX18* |
| 3 | SP1 | Sp1 transcription factor | 0.00127 | *CDH3, F10, PLAT, PTN, F2R* |
| 4 | TWIST1 | twist basic helix-loop-helix transcription factor 1 | 0.0017 | *F2R, MMP1* |
| 5 | MYB | v-myb myeloblastosis viral oncogene homolog (avian) | 0.0019 | *NMU, CTSG* |
| 6 | GATA3 | GATA binding protein 3 | 0.002 | *CDH3, MMP1* |
| 7 | POU2F1 | POU class 2 homeobox 1 | 0.002 | *GNRH1, SDC4* |
| 8 | JUN | jun proto-oncogene | 0.00221 | *PLAT, PTN, MMP1* |
| 9 | PPARG | peroxisome proliferator-activated receptor gamma | 0.00593 | *MMP1, CLDN2* |
| 10 | HDAC1 | histone deacetylase 1 | 0.00684 | *TJP1, CLDN7* |
| 11 | STAT1 | signal transducer and activator of transcription 1, 91kDa | 0.00945 | *XAF1, VIP* |
| 12 | CREB1 | cAMP responsive element binding protein 1 | 0.0108 | *VIP, PLAT* |
| 13 | STAT3 | signal transducer and activator of transcription 3 (acute-phase response factor) | 0.0255 | *F2R, MMP1* |
| 14 | RELA | v-rel reticuloendotheliosis viral oncogene homolog A (avian) | 0.097 | *SDC1, MMP1* |
| 15 | NFKB1 | nuclear factor of kappa light polypeptide gene enhancer in B-cells 1 | 0.0981 | *SDC1, MMP1* |

Using miRTarBase, microRNAs were identified as listed in Table 5.

**Table 5:** miRTarBase scanned microRNAs with respect to key regulators.

| S.N. | microRNA | Target Gene | Number of interactions | FDR | p-value |
| --- | --- | --- | --- | --- | --- |
| 1 | hsa-miR-1-3p | *RNF213, F2RL1, AP1S3* | 4 | 0.619 | 0.0885 |
| 2 | hsa-miR-335-5p | *F10, RNF213, VIP, CDH3, PLAT, CLDN7, XAF1, RASIP1, SOX18* | 9 | 0.619 | 0.043 |
| 3 | hsa-miR-526b-3p | *MMP1, F2RL1, F2R* | 3 | 0.619 | 0.114 |
| 4 | hsa-miR-615-3p | *TJP1, DNM2, AP2B1* | 3 | 0.619 | 0.229 |
| 5 | hsa-miR-106b-5p | *AP2B1, F2R, F2RL1* | 3 | 0.626 | 0.331 |
| 6 | hsa-miR-17-5p | *F2RL1, F2R, TJP1* | 3 | 0.641 | 0.378 |
| 7 | hsa-miR-124-3p | *DNM2, SDC4, RASIP1* | 3 | 0.666 | 0.513 |

## Discussion

The scarcity of knowledge regarding the genetic origins of Pancreatic Ductal Adenocarcinoma (PDAC) necessitates further investigation into its genetic aspects. Despite the identification of several candidate genes for PDAC in recent years, the underlying mechanism responsible for PDAC development remains unclear. To the best of our knowledge, our in-silico study constitutes the initial endeavor to investigate the Protein-Protein Interaction (PPI) network of PDAC and to explore the varying contributions of the proteins encoded by the candidate genes in regulating the entire network through topological analysis.

Our investigation of the topological properties of the initial PDAC network, comprising 605 nodes and 2698 edges, reveals a weak hierarchical and scale-free fractal structure. The Girvan Newman algorithm helps us to identify a two-tier organization of the network, with one level representing local clustering of mostly low-degree nodes into well-defined successive communities or modules, and the other level representing more global connectivity in which hubs serve as higher-order communication points between interconnected communities. The fractal state of the network signifies self-similar organization, while the scale-free nature contributes to network stability. These topological properties facilitate efficient information processing within the network.

The objective of predicting candidate genes for diseases, such as exploring the role of gene interactions, is a fundamental goal of the medical sciences and is crucial for effective treatment. To achieve this, it is necessary to conduct preliminary analysis to identify potential biomarkers. In this study, we applied differential expression analysis on RNA-seq data of PDAC to identify transcriptomic signatures that are characteristic of the disease, and then performed network analysis to better understand the interactions between genes. The results of this study can be used to identify potential biomarkers for PDAC and contribute to the field of pharmacogenomics, which has significant applications in drug discovery. We focused on network-regulated genes in this study and found that the network of classified genes from PDAC displays hierarchical characteristics, indicating that the network is organized at the sub-module level. This hierarchical nature of the network is important for understanding the functional regulation of the disease. Individual gene activities are less important in this process. Our networks, which comprise both up- and down-regulated genes, have led to the identification of 33 crucial regulators, including (*F10^↓^, PLAT^↑^,F2R^↑^, MMP1^↑^,CTSG^↓^, F2RL1^↓^,PDGFA^↓^, FGF13^↓^, PTN^↓^,APOC2^↑^, SDC1^↑^,SDC4^↑^, AP2B1^↑^, AP1S3^↑^, DNM2^↑^, EGFL7^↓^, RASIP1^↓^, SOX18^↓^, CLDN2^↑^, CLDN7^↑^, CDH3^↑^, CLDN5^↓^, TJP1^↑^, PCDH1^↓^, RNF213^↑^, EPSTI1^↑^, XAF1^↑^, VIP^↓^, GAL^↓^, GNRH1^↓^, NMU^↑^,VIPR2^↓^, and GNG11^↓^*). These regulators were identified through the analysis of motifs and module regulation, and their biological importance, roles in network activities and associated regulations, and potential as targets for disease have been established. Additionally, the biological activities and pathways in which these key regulator genes are involved have been identified.

Studies have shown the strong potential of F10 to improve treatment outcomes in acute myeloid leukemia, acute lymphocytic leukemia, glioblastoma, and prostate cancer (49). The PLAT gene has been studied in the context of cancer in several papers. PLAT, also known as tissue-type plasminogen activator, has been found to play a role in gefitinib resistance in non-small cell lung cancer (NSCLC) (50). Mutations and alterations in the FGFR2 gene have been found to play a significant role in the development and progression of various types of cancer, including endometrial cancer, breast cancer, and gastrointestinal/genitourinary tract cancers. These alterations include somatic hotspot mutations, structural amplifications, and fusions (51). Pleiotrophin (PTN) is a gene that has been found to be differentially expressed in various types of cancer, including hepatocellular carcinoma (HCC) (52), oral squamous cell carcinoma (OSCC), ovarian cancer, and breast cancer (BrCa). Tight junction proteins ZO-1, TJP1, TJP2, and TJP3 are scaffolding proteins that connect trans-membrane proteins like claudins and occludin to the actin cytoskeleton. They play a crucial role in maintaining the integrity of tight junctions and regulating para-cellular permeability (53). F2RL1, also known as Fc receptor-like 2, has been studied in the context of cancer. In metastatic breast cancer, decreased expression of FCRL2 mRNA was observed in brain metastatic tissues compared to primary breast tumors, and its expression in primary tumors was correlated with patient survival (54). PDGFA is a protein that has been implicated in cancer initiation and progression. It is up-regulated in several cancers, including colorectal cancer (CRC) (55). FGF13 has been found to play a role in cancer progression and treatment resistance. It has been shown to be associated with tumor growth and metastasis in pancreatic cancer (56). In human pluripotent stem cells, disrupting TJP1 leads to the activation of bone morphogenic protein-4 (BMP4) signaling and loss of patterning phenotype (57). CTSG has also been implicated in triple-negative breast cancer (TNBC), where it is overexpressed and correlated with a poor prognosis. CTGF, a protein that binds to CTSG, activates the FAK/Src/NF-κB p65 signaling axis, resulting in the up-regulation of Glut3 and enhanced aerobic glycolysis in TNBC cells (58). APOC2 has been identified as a potential diagnostic biomarker for cancer detection and as an auxiliary prognostic marker or marker for immunotherapy in certain tumor types (59). SDC1 has been shown to play a tumor-suppressor role in CRCs (60). In cervical cancer, SDC1 expression is associated with low differentiation and increased lymph node metastases (61). High SDC1 expression in cervical cancer is also correlated with a poor prognosis (62). The AP2B1 gene has been studied in various types of cancer. In lung cancer, the transcription factors AP2A and AP2B were found to promote the expression of the USP22 gene, which is associated with aggressive growth and therapy resistance (63). Patients with higher AP1S1 expression have higher estrogen receptor gene expression, increased risk of distant metastasis and lymph node metastasis, and worse overall survival rates (64). DNM2 appears to be a promising molecular target for the development of anti-invasive agents and has shown potential in reducing cell proliferation and inducing apoptosis in cancer cells (65).

EGFL7 is a gene that has been found to play a role in cancer. It has been identified as a driver gene for resistance to EGFR kinase inhibition in lung cancer cells (66).]. In addition, RASIP1 is negatively regulated by fork-head box O3 (FOXO3), which suppresses DLBCL cell proliferation. FOXO3 binds to the promoter sequence of RASIP1 and inhibits its transcription (67). In lung cancer, the inhibition of SOX18 with a specific inhibitor called Sm4 has shown cytotoxic effects on non-small cell lung cancer (NSCLC) cell lines, leading to cell cycle disruption and up-regulation of p21, a key regulator of cell cycle progression (68). In various cancers, including ovarian, testicular, endocervical, liver, and lung adenocarcinoma, CLDN7 is highly expressed and activates multiple signaling pathways involved in tumor growth, migration, invasion, and chemo-resistance (69). In OSCC, CDH3 is up-regulated and associated with a poor prognosis, promoting migration, invasion, and chemo-resistance in oral squamous cell carcinoma (70). CLDN5 expression levels differ significantly between cancer and normal tissues, and it has been confirmed in multiple studies (71). CLDN5 is implicated in the oncogenesis of diverse cancer types, highlighting its potential significance in cancer biology (72). PCDH1 enhances p65 nuclear localization by interacting with KPNB1, activating the NF-κB signaling pathway and promoting PDAC progression (73). PCDH1 can be used as a negative prognostic marker and a potential therapeutic target for PDAC patients (74). In breast cancer, RNF213 is differentially expressed in primary tumors and is correlated with overall survival in patients with basal-like sub-type breast cancer (75). Additionally, RNF213 knockdown disrupts angiogenesis and sensitizes endothelial cells to inflammation, leading to altered angiogenesis and potential links to Moyamoya disease. VIP expression has been associated with the transcription factor ZEB1, which is known to regulate EMT (76). In breast cancer, VIP receptor 2 (VIPR2) has been shown to promote cell proliferation and tumor growth through the cAMP/PKA/ERK signaling pathway (77). EPSTI1 interacts with valosin-containing protein to activate nuclear factor κ-light-chain-enhancer of activated B cells (NF-κB) and inhibit apoptosis (78). EPSTI1 has also been implicated in immune response, as it promotes the expression of viral response genes and is associated with immune privilege and autoimmune diseases (79). NMU is expressed at higher levels in tumor tissues compared to normal tissues, and its expression has been associated with poor prognosis and shorter overall survival in cancer patients (80). Studies have shown that VIPR2 overexpression promotes cell proliferation in breast cancer cell lines and exacerbates tumor growth in vivo (77). Furthermore, VIPR2 has been shown to form homodimers and oligomers, which are involved in VIP-induced cancer cell migration (81). The expression of GNG11 mRNA is down-regulated in ovarian cancer patients, and its high expression is associated with poor prognosis (82). GNG11 may play a crucial role in the biological process of ovarian cancer through the ECM-receptor interaction pathway (83).

We further performed the pathway crosstalk analysis to explore the interactions among significantly enriched pathways. Genes VIPR2, GAL, F2R, NMU, F2RL1, GNRH1, CTSG, VIP were involved in neuroactive ligand-receptor interaction pathways. Key regulator genes CLDN5, CDH3, SDC4, CLDN7, SDC1, CLDN2 were active in Cell adhesion molecules and CLDN5, CLDN7, CLDN2 enriched in Leukocyte trans-endothelial migration pathway. Go classification of key signature genes indicates the regulation of positive regulation of cell proliferation, positive regulation of protein kinase B signaling, calcium-independent cell-cell adhesion via plasma membrane cell-adhesion molecules related biological process. While important cellular process are extracellular region, plasma membrane, and cell surface. Key regulator genes were enriched in G-protein beta-sub-unit binding, receptor binding, serine-type endopeptidase activity and growth factor activity as molecular function as listed in Table 6.

**Table 6.** Enrichment analysis results for key signature genes.

| Category | Description | Genes | P-value | Fold Enrichment |
| --- | --- | --- | --- | --- |
| MF | G-protein beta-subunit binding | *F2R, F2RL1, GNG11* | 0.000697 | 74.125 |
| MF | receptor binding | *EGFL7, F2R, NMU, F2RL1, PLAT* | 0.00433306 | 7.196601942 |
| MF | serine-type endopeptidase activity | *F10, MMP1, CTSG, PLAT* | 0.003455303 | 12.68449198 |
| MF | growth factor activity | *PDGFA, PTN, FGF13* | 0.040571326 | 9.076530612 |
| CC | extracellular region | *EGFL7, F10, MMP1, F2R, PDGFA, PLAT, PTN, GAL, APOC2, NMU, GNRH1, CTSG, VIP, FGF13* | 0.00000756 | 4.084387709 |
| CC | plasma membrane | *VIPR2, F10, SDC4, F2R, AP2B1, PTN, CLDN2, GNG11, DNM2, TJP1, CLDN5, CDH3, CLDN7, F2RL1, SDC1, CTSG, PCDH1, FGF13* | 0.000728 | 2.129213998 |
| CC | cell surface | *EGFL7, SDC4, F2R, PDGFA, SDC1, CTSG, PLAT* | 0.00038 | 6.868351064 |
| BP | positive regulation of cell proliferation | *TJP1, CLDN5, F2R, CLDN7, PDGFA, PTN, VIP* | 0.00019 | 7.811582569 |
| BP | positive regulation of protein kinase B signaling | *F10, F2R, PDGFA, F2RL1* | 0.003123059 | 13.15 |
| BP | calcium-independent cell-cell adhesion via plasma membrane cell-adhesion molecules | *CLDN5, CLDN7, CLDN2* | 0.000506 | 86.88392857 |
| KEGG | Neuroactive ligand-receptor interaction | *VIPR2, GAL, F2R, NMU, F2RL1, GNRH1, CTSG, VIP* | 0.0000453 | 7.486430518 |
| KEGG | Cell adhesion molecules | *CLDN5, CDH3, SDC4, CLDN7, SDC1, CLDN2* | 0.0000634 | 13.04202532 |

Further, gene-drug interaction for regulator genes retrieved drugs against 14 genes as listed in Table 3.

Transcription factor screening for key regulator genes inferred 15 transcription factors as listed in Table 4. Some well-known transcription factors among them are STAT1-signal transducer and activator of transcription 1, STAT3-signal transducer and activator of transcription 3 (acute-phase response factor), CREB1-cAMP responsive element binding protein 1, RELA-v-rel reticuloendotheliosis viral oncogene homolog A (avian),and NFKB1-nuclear factor of kappa light polypeptide gene enhancer in B-cells 1.

Next, using miRTarBase, 7 microRNAs namely hsa-miR-1-3p-RNF213, F2RL1, AP1S3; hsa-miR-335-5p-F10, RNF213, VIP, CDH3, PLAT, CLDN7, XAF1, RASIP1, SOX18; hsa-miR-526b-3p-MMP1, F2RL1, F2R; hsa-miR-615-3p-TJP1, DNM2, AP2B1; hsa-miR-106b-5p-AP2B1, F2R, F2RL1; hsa-miR-17-5p-F2RL1, F2R, TJP1; and hsa-miR-124-3p-DNM2, SDC4, RASIP1 were found and listed in Table 5.

In this study, we applied a systematic and extensible methodology, which finds significant key regulators in PDAC RNA-seq data. This study suggested that gene expression profiling can be used to differentiate and identify patients from their healthy counterparts. However, experimental validation of these results in large sample size would confirm the reliableness of these key regulators and facilitate the designing of an economical and susceptive molecular diagnostic.

### Limitation of the Study

Nevertheless, further investigation is necessary to verify the expression and function of the key regulators identified in PDAC, given the constrained sample size.

## Conclusion

We conducted a comprehensive analysis using one RNA-seq gene expression profile, comparing individuals with PDAC to healthy controls. Our aim was to identify Differentially Expressed Genes (DEGs) and elucidate their biological functions through pathway enrichment analysis. Additionally, we explored the topological features of the gene interaction network, uncovering significant key regulators associated with PDAC. Moreover, we investigated approved drugs, transcription factors and mi-RNAs targeting these key regulators. These genes are anticipated to play a crucial role in PDAC progression. This study can be considered as very useful in the context of personalized treatment plan for PDAC patients. Personalized treatment in cancer represents a paradigm shift towards individualized and precision-based approaches to diagnosis, prognosis, and therapy selection, with the ultimate goal of improving patient outcomes and quality of life.

## Acknowledgments

KB is thankful to the Indian Council of Medical Research (ICMR) (BMI/11(92)/2022) for providing the research fellowship. We are thankful to the HPC of Ashoka University for providing the necessary infrastructure to help us successfully conduct our research work. We express our gratitude to Dr. Ram Nayan Verma for his valuable and insightful contributions to the discussions on the network analysis method utilized in this study.

## Conflict of Interest

All authors declare no conflicts of interest.

## Notes

### Competing Interest Statement

The authors have declared no competing interest.

## References

1. Calimano-Ramirez LF, Daoud T, Gopireddy DR, Morani AC, Waters R, Gumus K, et al. Pancreatic acinar cell carcinoma: A comprehensive review. World J Gastroenterol [Internet]. 2022 Oct 21 [cited 2024 Mar 10];28(40):5827–44. Available from: https://www.wjgnet.com/1007-9327/full/v28/i40/5827.htm

2. Anderson EM, Thomassian S, Gong J, Hendifar A, Osipov A. Advances in Pancreatic Ductal Adenocarcinoma Treatment. Cancers [Internet]. 2021 Nov 3 [cited 2024 Mar 10];13(21):5510. Available from: https://www.mdpi.com/2072-6694/13/21/5510

3. Tran K, Prall OWJ, Mitchell C, Chou A, Gill AJ, Grimmond SM, et al. Diagnosing primary pancreatic acinar cell carcinoma – Clinical correlation of radiological/molecular imaging, histopathologic features and whole genome/transcriptome profiling, and review of the literature. Current Problems in Cancer: Case Reports [Internet]. 2023 Mar [cited 2024 Mar 10];9:100213. Available from: https://linkinghub.elsevier.com/retrieve/pii/S2666621922000771

4. Elsayed M, Abdelrahim M. The Latest Advancement in Pancreatic Ductal Adenocarcinoma Therapy: A Review Article for the Latest Guidelines and Novel Therapies. Biomedicines [Internet]. 2021 Apr 6 [cited 2024 Mar 10];9(4):389. Available from: https://www.mdpi.com/2227-9059/9/4/389

5. Mere Del Aguila E, Tang XH, Gudas LJ. Pancreatic Ductal Adenocarcinoma: New Insights into the Actions of Vitamin A. Oncol Res Treat [Internet]. 2022 [cited 2024 Mar 10];45(5):291–8. Available from: https://karger.com/doi/10.1159/000522425

6. Taherian M, Wang H, Wang H. Pancreatic Ductal Adenocarcinoma: Molecular Pathology and Predictive Biomarkers. Cells [Internet]. 2022 Sep 29 [cited 2024 Mar 10];11(19):3068. Available from: https://www.mdpi.com/2073-4409/11/19/3068

7. Sakaguchi R, Yamada R, Nose K, Tanaka T, Murashima Y, Tsuboi J, et al. Pancreatic Mixed Acinar-neuroendocrine-ductal Carcinoma: A Case Report and Literature Review. Intern Med [Internet]. 2023 Nov 15 [cited 2024 Mar 10];62(22):3347–53. Available from: https://www.jstage.jst.go.jp/article/internalmedicine/62/22/62_1298-22/_article

8. Alzhrani R, Alsaab HO, Vanamala K, Bhise K, Tatiparti K, Barari A, et al. Overcoming the Tumor Microenvironmental Barriers of Pancreatic Ductal Adenocarcinomas for Achieving Better Treatment Outcomes. Advanced Therapeutics [Internet]. 2021 Jun [cited 2024 Mar 10];4(6):2000262. Available from: https://onlinelibrary.wiley.com/doi/10.1002/adtp.202000262

9. Kang E, Choi YS, Oh HC, Do JH, Hong SU, Lee SE. Rapidly Growing Acinar Cell Carcinoma of the Pancreatic Head: A Case Report and Literature Review. Korean J Pancreas Biliary Tract [Internet]. 2022 Jan 31 [cited 2024 Mar 10];27(1):54–9. Available from: http://kjpbt.org/journal/view.php?doi=10.15279/kpba.2022.27.1.54

10. Elsayed M, Abdelrahim M. The Latest Advancement in Pancreatic Ductal Adenocarcinoma Therapy: A Review Article for the Latest Guidelines and Novel Therapies. Biomedicines [Internet]. 2021 Apr 6 [cited 2024 Mar 10];9(4):389. Available from: https://www.mdpi.com/2227-9059/9/4/389

11. Skorupan N, Palestino Dominguez M, Ricci SL, Alewine C. Clinical Strategies Targeting the Tumor Microenvironment of Pancreatic Ductal Adenocarcinoma. Cancers [Internet]. 2022 Aug 30 [cited 2024 Mar 10];14(17):4209. Available from: https://www.mdpi.com/2072-6694/14/17/4209

12. Truong LH, Pauklin S. Pancreatic Cancer Microenvironment and Cellular Composition: Current Understandings and Therapeutic Approaches. Cancers [Internet]. 2021 Oct 8 [cited 2024 Mar 10];13(19):5028. Available from: https://www.mdpi.com/2072-6694/13/19/5028

13. Sally Á, McGowan R, Finn K, Moran BM. Current and Future Therapies for Pancreatic Ductal Adenocarcinoma. Cancers [Internet]. 2022 May 13 [cited 2024 Mar 10];14(10):2417. Available from: https://www.mdpi.com/2072-6694/14/10/2417

14. Chakrabarti S, Kamgar M, Mahipal A. Systemic Therapy of Metastatic Pancreatic Adenocarcinoma: Current Status, Challenges, and Opportunities. Cancers [Internet]. 2022 May 24 [cited 2024 Mar 10];14(11):2588. Available from: https://www.mdpi.com/2072-6694/14/11/2588

15. Sarfraz H, Saha A, Jhaveri K, Kim DW. Review of Current Systemic Therapy and Novel Systemic Therapy for Pancreatic Ductal Adenocarcinoma. Current Oncology [Internet]. 2023 May 26 [cited 2024 Mar 10];30(6):5322–36. Available from: https://www.mdpi.com/1718-7729/30/6/404

16. Dhaliwal J. Exploring the Therapeutic Opportunities of the Tumour Microenvironment in Treating Pancreatic Ductal Adenocarcinoma: A Literature Review. URNCST Journal [Internet]. 2022 Nov 15 [cited 2024 Mar 10];6(11):1–9. Available from: https://urncst.com/index.php/urncst/article/view/406

17. Biagi JJ, Cosby R, Bahl M, Elfiki T, Goodwin R, Hallet J, et al. Adjuvant Chemotherapy and Radiotherapy in Resected Pancreatic Ductal Adenocarcinoma: A Systematic Review and Clinical Practice Guideline. Current Oncology [Internet]. 2023 Jul 8 [cited 2024 Mar 10];30(7):6575–86. Available from: https://www.mdpi.com/1718-7729/30/7/482

18. Afghani E, Klein AP. Pancreatic Adenocarcinoma. Hematology/Oncology Clinics of North America [Internet]. 2022 Oct [cited 2024 Mar 10];36(5):879–95. Available from: https://linkinghub.elsevier.com/retrieve/pii/S0889858822000703

19. Zhao F, Yang D, Xu T, He J, Guo J, Li X. New treatment insights into pancreatic acinar cell carcinoma: case report and literature review. Front Oncol [Internet]. 2023 Jul 3 [cited 2024 Mar 10];13:1210064. Available from: https://www.frontiersin.org/articles/10.3389/fonc.2023.1210064/full

20. Barabási AL, Gulbahce N, Loscalzo J. Network medicine: a network-based approach to human disease. Nat Rev Genet [Internet]. 2011 Jan [cited 2024 Mar 10];12(1):56–68. Available from: https://www.nature.com/articles/nrg2918

21. Chen SJ, Liao DL, Chen CH, Wang TY, Chen KC. Construction and Analysis of Protein-Protein Interaction Network of Heroin Use Disorder. Sci Rep [Internet]. 2019 Mar 21 [cited 2024 Mar 10];9(1):4980. Available from: https://www.nature.com/articles/s41598-019-41552-z

22. Verma RN, Malik MdZ, Subbarao N, Singh GP, Sinha DN. Entamoeba histolytica HM-1: IMSS gene expression profiling identifies key hub genes, potential biomarkers, and pathways in Amoebiasis infection: a systematic network meta-analysis. Bioscience Reports [Internet]. 2022 Oct 28 [cited 2024 Mar 10];42(10):BSR20220191. Available from: https://portlandpress.com/bioscirep/article/42/10/BSR20220191/231689/Entamoeba-histolytica-HM-1-IMSS-gene-expression

23. Browne F, Wang H, Zheng H. Investigating the impact human protein–protein interaction networks have on disease-gene analysis. Int J Mach Learn & Cyber [Internet]. 2018 Mar [cited 2024 Mar 10];9(3):455–64. Available from: http://link.springer.com/10.1007/s13042-016-0503-5

24. Albert R, Barabási AL. Statistical mechanics of complex networks. Rev Mod Phys [Internet]. 2002 Jan 30 [cited 2024 Mar 10];74(1):47–97. Available from: https://link.aps.org/doi/10.1103/RevModPhys.74.47

25. Verma RN, Malik MdZ, Singh GP, Subbarao N. Identification of key proteins in host–pathogen interactions between Mycobacterium tuberculosis and Homo sapiens: A systematic network theoretical approach. Healthcare Analytics [Internet]. 2022 Nov [cited 2024 Mar 10];2:100052. Available from: https://linkinghub.elsevier.com/retrieve/pii/S2772442522000193

26. Rajamani D, Bhasin MK. Identification of key regulators of pancreatic cancer progression through multidimensional systems-level analysis. Genome Med [Internet]. 2016 Dec [cited 2024 Mar 10];8(1):38. Available from: http://genomemedicine.biomedcentral.com/articles/10.1186/s13073-016-0282-3

27. Liu J, Xiong Q, Shi W, Shi X, Wang K. Evaluating the importance of nodes in complex networks. Physica A: Statistical Mechanics and its Applications [Internet]. 2016 Jun [cited 2024 Mar 10];452:209–19. Available from: https://linkinghub.elsevier.com/retrieve/pii/S0378437116002156

28. Wu H, Tian W, Tai X, Li X, Li Z, Shui J, et al. Identification and functional analysis of novel oncogene DDX60L in pancreatic ductal adenocarcinoma. BMC Genomics. 2021 Nov 18;22(1):833.

29. Dembélé D, Kastner P. Fold change rank ordering statistics: a new method for detecting differentially expressed genes. BMC Bioinformatics [Internet]. 2014 Dec [cited 2024 Mar 10];15(1):14. Available from: https://bmcbioinformatics.biomedcentral.com/articles/10.1186/1471-2105-15-14

30. Ritchie ME, Phipson B, Wu D, Hu Y, Law CW, Shi W, et al. limma powers differential expression analyses for RNA-sequencing and microarray studies. Nucleic Acids Research [Internet]. 2015 Apr 20 [cited 2024 Mar 10];43(7):e47–e47. Available from: http://academic.oup.com/nar/article/43/7/e47/2414268/limma-powers-differential-expression-analyses-for

31. Baltagi BH, Li D. LM Tests for Functional Form and Spatial Error Correlation. International Regional Science Review [Internet]. 2001 Apr [cited 2024 Mar 10];24(2):194–225. Available from: http://journals.sagepub.com/doi/10.1177/016001760102400202

32. Datta S, Datta S. Empirical Bayes screening of many p-values with applications to microarray studies. Bioinformatics [Internet]. 2005 May 1 [cited 2024 Mar 10];21(9):1987–94. Available from: https://academic.oup.com/bioinformatics/article-lookup/doi/10.1093/bioinformatics/bti301

33. Duncan DB. Multiple Range and Multiple F Tests. Biometrics [Internet]. 1955 Mar [cited 2024 Mar 10];11(1):1. Available from: https://www.jstor.org/stable/3001478?origin=crossref

34. Lakens D. Calculating and reporting effect sizes to facilitate cumulative science: a practical primer for t-tests and ANOVAs. Front Psychol [Internet]. 2013 [cited 2024 Mar 10];4. Available from: http://journal.frontiersin.org/article/10.3389/fpsyg.2013.00863/abstract

35. Wilkinson L. ggplot2: Elegant Graphics for Data Analysis by WICKHAM, H. Biometrics [Internet]. 2011 Jun [cited 2024 Mar 10];67(2):678–9. Available from: https://academic.oup.com/biometrics/article/67/2/678-679/7381027

36. Shannon P, Markiel A, Ozier O, Baliga NS, Wang JT, Ramage D, et al. Cytoscape: A Software Environment for Integrated Models of Biomolecular Interaction Networks. Genome Res [Internet]. 2003 Nov [cited 2024 Mar 10];13(11):2498–504. Available from: http://genome.cshlp.org/lookup/doi/10.1101/gr.1239303

37. Ashburner M, Ball CA, Blake JA, Botstein D, Butler H, Cherry JM, et al. Gene Ontology: tool for the unification of biology. Nat Genet [Internet]. 2000 May [cited 2024 Mar 10];25(1):25–9. Available from: https://www.nature.com/articles/ng0500_25

38. Barabási AL, Oltvai ZN. Network biology: understanding the cell’s functional organization. Nat Rev Genet [Internet]. 2004 Feb [cited 2024 Mar 10];5(2):101–13. Available from: https://www.nature.com/articles/nrg1272

39. Albert R, Barabási AL. Statistical mechanics of complex networks. Rev Mod Phys [Internet]. 2002 Jan 30 [cited 2024 Mar 10];74(1):47–97. Available from: https://link.aps.org/doi/10.1103/RevModPhys.74.47

40. Csárdi G, Nepusz T, Müller K, Horvát S, Traag V, Zanini F, et al. igraph for R: R interface of the igraph library for graph theory and network analysis [Internet]. [object Object]; 2024 [cited 2024 Mar 10]. Available from: https://zenodo.org/doi/10.5281/zenodo.7682609

41. Sadi S, Oguducu S, Uyar AS. An efficient community detection method using parallel clique-finding ants. In: IEEE Congress on Evolutionary Computation [Internet]. Barcelona, Spain: IEEE; 2010 [cited 2024 Mar 10]. p. 1–7. Available from: http://ieeexplore.ieee.org/document/5586496/

42. Licursi V, Conte F, Fiscon G, Paci P. MIENTURNET: an interactive web tool for microRNA-target enrichment and network-based analysis. BMC Bioinformatics [Internet]. 2019 Dec [cited 2024 Mar 10];20(1):545. Available from: https://bmcbioinformatics.biomedcentral.com/articles/10.1186/s12859-019-3105-x

43. Hsu SD, Lin FM, Wu WY, Liang C, Huang WC, Chan WL, et al. miRTarBase: a database curates experimentally validated microRNA–target interactions. Nucleic Acids Research [Internet]. 2011 Jan 1 [cited 2024 Mar 10];39(suppl_1):D163–9. Available from: https://academic.oup.com/nar/article-lookup/doi/10.1093/nar/gkq1107

44. Cotto KC, Wagner AH, Feng YY, Kiwala S, Coffman AC, Spies G, et al. DGIdb 3.0: a redesign and expansion of the drug–gene interaction database. Nucleic Acids Research [Internet]. 2018 Jan 4 [cited 2024 Mar 10];46(D1):D1068–73. Available from: https://academic.oup.com/nar/article/46/D1/D1068/4634012

45. Clauset A, Shalizi CR, Newman MEJ. Power-Law Distributions in Empirical Data. SIAM Rev [Internet]. 2009 Nov 4 [cited 2024 Mar 10];51(4):661–703. Available from: http://epubs.siam.org/doi/10.1137/070710111

46. Rajamani D, Bhasin MK. Identification of key regulators of pancreatic cancer progression through multidimensional systems-level analysis. Genome Medicine [Internet]. 2016 May 3 [cited 2019 Dec 4];8(1):38. Available from: 10.1186/s13073-016-0282-3

47. Lawyer G. Understanding the influence of all nodes in a network. Scientific Reports [Internet]. 2015 Mar 2 [cited 2019 Dec 4];5(1):1–9. Available from: https://www.nature.com/articles/srep08665

48. Liu J, Xiong Q, Shi W, Shi X, Wang K. Evaluating the importance of nodes in complex networks. Physica A: Statistical Mechanics and its Applications [Internet]. 2016 Jun 15 [cited 2019 Dec 4];452:209–19. Available from: http://www.sciencedirect.com/science/article/pii/S0378437116002156

49. Potez M, Trappetti V, Bouchet A, Fernandez-Palomo C, Güç E, Kilarski WW, et al. Characterization of a B16-F10 melanoma model locally implanted into the ear pinnae of C57BL/6 mice. Mattei F, editor. PLoS ONE [Internet]. 2018 Nov 5 [cited 2024 Mar 10];13(11):e0206693. Available from: https://dx.plos.org/10.1371/journal.pone.0206693

50. Martínez-Puente DH, Pérez-Trujillo JJ, Zavala-Flores LM, García-García A, Villanueva-Olivo A, Rodríguez-Rocha H, et al. Plasmid DNA for Therapeutic Applications in Cancer. Pharmaceutics [Internet]. 2022 Sep 3 [cited 2024 Mar 10];14(9):1861. Available from: https://www.mdpi.com/1999-4923/14/9/1861

51. Dixit G, Gonzalez-Bosquet J, Skurski J, Devor EJ, Dickerson EB, Nothnick WB, et al. FGFR2 mutations promote endometrial cancer progression through dual engagement of EGFR and Notch signalling pathways. Clinical & Translational Med [Internet]. 2023 May [cited 2024 Mar 10];13(5):e1223. Available from: https://onlinelibrary.wiley.com/doi/10.1002/ctm2.1223

52. Lin C, Chen Y, Zhang F, Zhu P, Yu L, Chen W. Single-cell RNA sequencing reveals the mediatory role of cancer-associated fibroblast PTN in hepatitis B virus cirrhosis-HCC progression. Gut Pathog [Internet]. 2023 May 31 [cited 2024 Mar 10];15(1):26. Available from: https://gutpathogens.biomedcentral.com/articles/10.1186/s13099-023-00554-z

53. Chan S, Wang X, Wang Z, Du Y, Zuo X, Chen J, et al. CTSG Suppresses Colorectal Cancer Progression through Negative Regulation of Akt/mTOR/Bcl2 Signaling Pathway. Int J Biol Sci [Internet]. 2023 [cited 2024 Mar 10];19(7):2220–33. Available from: https://www.ijbs.com/v19p2220.htm

54. Mamoor S. FCRL2 is a differentially expressed gene in brain metastatic human breast cancer. [Internet]. Open Science Framework; 2023 Jun [cited 2024 Mar 10]. Available from: https://osf.io/tudkg

55. Li J, Zhi X, Sun Y, Chen M, Yao L. The PDGF Family Is Associated with Activated Tumor Stroma and Poor Prognosis in Ovarian Cancer. Sun Y, editor. Disease Markers [Internet]. 2022 Sep 26 [cited 2024 Mar 10];2022:1–19. Available from: https://www.hindawi.com/journals/dm/2022/5940049/

56. Sun Y, Li D, Liu H, Huang Y, Meng F, Tang J, et al. PHF13 epigenetically activates TGFβ driven epithelial to mesenchymal transition. Cell Death Dis [Internet]. 2022 May 21 [cited 2024 Mar 10];13(5):487. Available from: https://www.nature.com/articles/s41419-022-04940-4

57. Vasic I, Libby ARG, Maslan A, Bulger EA, Zalazar D, Krakora Compagno MZ, et al. Loss of TJP1 disrupts gastrulation patterning and increases differentiation toward the germ cell lineage in human pluripotent stem cells. Developmental Cell [Internet]. 2023 Aug [cited 2024 Mar 10];58(16):1477–1488.e5. Available from: https://linkinghub.elsevier.com/retrieve/pii/S1534580723002654

58. Huang D, Wang X, Liu Y, Huang Z, Hu X, Hu W, et al. Multi-omic analysis suggests tumor suppressor genes evolved specific promoter features to optimize cancer resistance. Briefings in Bioinformatics [Internet]. 2021 Sep 2 [cited 2024 Mar 10];22(5):bbab040. Available from: https://academic.oup.com/bib/article/doi/10.1093/bib/bbab040/6200210

59. Deng H, Li J, Shah AA, Ge L, Ouyang W. Comprehensive in-silico analysis of deleterious SNPs in APOC2 and APOA5 and their differential expression in cancer and cardiovascular diseases conditions. Genomics [Internet]. 2023 Mar [cited 2024 Mar 10];115(2):110567. Available from: https://linkinghub.elsevier.com/retrieve/pii/S0888754323000113

60. Yang Z, Chen S, Ying H, Yao W. Targeting syndecan-1: new opportunities in cancer therapy. American Journal of Physiology-Cell Physiology [Internet]. 2022 Jul 1 [cited 2024 Mar 10];323(1):C29–45. Available from: https://journals.physiology.org/doi/10.1152/ajpcell.00024.2022

61. Hilgers K, Ibrahim SA, Kiesel L, Greve B, Espinoza-Sánchez NA, Götte M. Differential Impact of Membrane-Bound and Soluble Forms of the Prognostic Marker Syndecan-1 on the Invasiveness, Migration, Apoptosis, and Proliferation of Cervical Cancer Cells. Front Oncol [Internet]. 2022 Jan 27 [cited 2024 Mar 10];12:803899. Available from: https://www.frontiersin.org/articles/10.3389/fonc.2022.803899/full

62. Song G, Ma Y, Ma Y, Liu P, Hou L, Xu Z, et al. miR-335-5p Targets SDC1 to Regulate the Progression of Breast Cancer. Crit Rev Eukaryot Gene Expr [Internet]. 2022 [cited 2024 Mar 10];32(6):21–31. Available from: https://www.dl.begellhouse.com/journals/6dbf508d3b17c437,40058b3a3fcd229f,0c88200b528944ae.html

63. Sun T, Zhang K, Li W, Liu Y, Pangeni RP, Li A, et al. Transcription factor AP2 enhances malignancy of non-small cell lung cancer through upregulation of USP22 gene expression. Cell Commun Signal [Internet]. 2022 Sep 19 [cited 2024 Mar 10];20(1):147. Available from: https://biosignaling.biomedcentral.com/articles/10.1186/s12964-022-00946-9

64. Dastmalchi N, Akbarzadeh S, Amini F, Rajabi A, Safaralizadeh R. Alterations in the expression levels of long intergenic non-coding RNA APOC1P1-3 in cervical cancer tissue samples. Nucleosides, Nucleotides & Nucleic Acids [Internet]. 2023 Jun 3 [cited 2024 Mar 10];42(6):495–505. Available from: https://www.tandfonline.com/doi/full/10.1080/15257770.2022.2160459

65. Trochet D, Bitoun M. A review of Dynamin 2 involvement in cancers highlights a promising therapeutic target. J Exp Clin Cancer Res [Internet]. 2021 Dec [cited 2024 Mar 10];40(1):238. Available from: https://jeccr.biomedcentral.com/articles/10.1186/s13046-021-02045-y

66. Wang Y, Chen P, Zhao M, Cao H, Zhao Y, Ji M, et al. EGFL7 drives the evolution of resistance to EGFR inhibitors in lung cancer by activating NOTCH signaling. Cell Death Dis [Internet]. 2022 Oct 29 [cited 2024 Mar 10];13(10):910. Available from: https://www.nature.com/articles/s41419-022-05354-y

67. Liu Y, Wu X, Feng Y, Jiang Q, Zhang S, Wang Q, et al. Insights into the Oncogenic, Prognostic, and Immunological Role of BRIP1 in Pan-Cancer: A Comprehensive Data-Mining-Based Study. Liu Y, editor. Journal of Oncology [Internet]. 2023 Apr 28 [cited 2024 Mar 10];2023:1–19. Available from: https://www.hindawi.com/journals/jo/2023/4104639/

68. Rodak O, Mrozowska M, Rusak A, Gomułkiewicz A, Piotrowska A, Olbromski M, et al. Targeting SOX18 Transcription Factor Activity by Small-Molecule Inhibitor Sm4 in Non-Small Lung Cancer Cell Lines. IJMS [Internet]. 2023 Jul 11 [cited 2024 Mar 10];24(14):11316. Available from: https://www.mdpi.com/1422-0067/24/14/11316

69. Hou Y, Hou L, Liang Y, Zhang Q, Hong X, Wang Y, et al. The p53-inducible CLDN7 regulates colorectal tumorigenesis and has prognostic significance. Neoplasia [Internet]. 2020 Nov [cited 2024 Mar 10];22(11):590–603. Available from: https://linkinghub.elsevier.com/retrieve/pii/S1476558620301469

70. Feng R, Chen Y, Liu Y, Zhou Q, Zhang W. The role of B7-H3 in tumors and its potential in clinical application. International Immunopharmacology [Internet]. 2021 Dec [cited 2024 Mar 10];101:108153. Available from: https://linkinghub.elsevier.com/retrieve/pii/S156757692100789X

71. Hashimoto Y, Greene C, Munnich A, Campbell M. The CLDN5 gene at the blood-brain barrier in health and disease. Fluids Barriers CNS [Internet]. 2023 Mar 28 [cited 2024 Mar 10];20(1):22. Available from: https://fluidsbarrierscns.biomedcentral.com/articles/10.1186/s12987-023-00424-5

72. Xu L, Shao F, Luo T, Li Q, Tan D, Tan Y. Pan-Cancer Analysis Identifies CHD5 as a Potential Biomarker for Glioma. IJMS [Internet]. 2022 Jul 30 [cited 2024 Mar 10];23(15):8489. Available from: https://www.mdpi.com/1422-0067/23/15/8489

73. Ye Z, Yang Y, Wei Y, Li L, Wang X, Zhang J. PCDH1 promotes progression of pancreatic ductal adenocarcinoma via activation of NF-κB signalling by interacting with KPNB1. Cell Death Dis [Internet]. 2022 Jul 21 [cited 2024 Mar 10];13(7):633. Available from: https://www.nature.com/articles/s41419-022-05087-y

74. Zheng Z, Luan N, Tu K, Liu F, Wang J, Sun J. The roles of protocadherin-7 in colorectal cancer cells on cell proliferation and its chemoresistance. Front Pharmacol [Internet]. 2023 Mar 30 [cited 2024 Mar 10];14:1072033. Available from: https://www.frontiersin.org/articles/10.3389/fphar.2023.1072033/full

75. Zhang L, Rashad S, Zhou Y, Niizuma K, Tominaga T. RNF213 loss of function reshapes vascular transcriptome and spliceosome leading to disrupted angiogenesis and aggravated vascular inflammatory responses. J Cereb Blood Flow Metab [Internet]. 2022 Nov [cited 2024 Mar 10];42(11):2107–22. Available from: http://journals.sagepub.com/doi/10.1177/0271678X221110679

76. Asano S, Ono A, Sakamoto K, Hayata-Takano A, Nakazawa T, Tanimoto K, et al. Vasoactive intestinal peptide receptor 2 signaling promotes breast cancer cell proliferation by enhancing the ERK pathway. Peptides [Internet]. 2023 Mar [cited 2024 Mar 10];161:170940. Available from: https://linkinghub.elsevier.com/retrieve/pii/S0196978123000025

77. Kittikulsuth W, Nakano D, Kitada K, Uyama T, Ueda N, Asano E, et al. Vasoactive intestinal peptide blockade suppresses tumor growth by regulating macrophage polarization and function in CT26 tumor-bearing mice. Sci Rep [Internet]. 2023 Jan 17 [cited 2024 Mar 10];13(1):927. Available from: https://www.nature.com/articles/s41598-023-28073-6

78. Kim YH, Lee JR, Hahn MJ. Regulation of inflammatory gene expression in macrophages by epithelial-stromal interaction 1 (Epsti1). Biochemical and Biophysical Research Communications [Internet]. 2018 Feb [cited 2024 Mar 10];496(2):778–83. Available from: https://linkinghub.elsevier.com/retrieve/pii/S0006291X17323987

79. Justin Gray And Jihe Zhao. Implications of Epithelial–Stromal Interaction 1 in Diseases Associated with Inflammatory Signaling. Cell Communication Insights [Internet]. 2016 Feb 29 [cited 2024 Mar 10];8:1–6. Available from: http://access.portico.org/stable?au=phw17wtv7c8

80. Qi X, Liu P, Wang Y, Xue J, An Y, Zhao C. Insights Into the Research Status of Neuromedin U: A Bibliometric and Visual Analysis From 1987 to 2021. Front Med [Internet]. 2022 Feb 22 [cited 2024 Mar 10];9:773000. Available from: https://www.frontiersin.org/articles/10.3389/fmed.2022.773000/full

81. Asano S, Ozasa K, Nakazawa T, Waschek A J, Ago Y. Functional importance of the oligomer formation of the neuropeptide receptor VIPR2. Proceedings for Annual Meeting of The Japanese Pharmacological Society [Internet]. 2022 [cited 2024 Mar 10];95(0):3-O-125. Available from: https://www.jstage.jst.go.jp/article/jpssuppl/95/0/95_3-O-125/_article/-char/ja/

82. Król W, Machelak W, Zielińska M. GDF11 as a friend or an enemy in the cancer biology? Biochimica et Biophysica Acta (BBA) - Reviews on Cancer [Internet]. 2023 Sep [cited 2024 Mar 10];1878(5):188944. Available from: https://linkinghub.elsevier.com/retrieve/pii/S0304419X23000938

83. Jiang MM, Zhao F, Lou TT. Assessment of Significant Pathway Signaling and Prognostic Value of GNG11 in Ovarian Serous Cystadenocarcinoma. IJGM [Internet]. 2021 Jun [cited 2024 Mar 10];Volume 14:2329–41. Available from: https://www.dovepress.com/assessment-of-significant-pathway-signaling-and-prognostic-value-of-gn-peer-reviewed-fulltext-article-IJGM

